# Interferon-γ Elicits Pathological Hallmarks of ALS in Human Motor Neurons

**DOI:** 10.1101/2022.11.18.517141

**Authors:** Changho Chun, Jung Hyun Lee, Alec S.T. Smith, David L. Mack, Mark Bothwell, Paul Nghiem

## Abstract

Neuroinflammation is an established factor contributing to amyotrophic lateral sclerosis (ALS) pathology, implicating the possible detrimental effects of inflammatory cytokines on motor neurons. The RNA/DNA-binding protein TDP-43 has emerged as a pivotal actor in ALS, because TDP-43 mutations cause familial ALS and loss of nuclear TDP-43, associated with its redistribution into cytoplasmic aggregates (TDP-43 proteinopathy) in motor neurons occurs in 97% of ALS cases. However, mechanisms linking neuroinflammation to TDP-43 mislocalization have not been described. Programmed death-ligand 1 (PD-L1) is an immune-modulatory protein, highly expressed on cell surfaces following acute inflammatory stress. To determine which inflammatory cytokines might impact motor neuron function, seven cytokines known to be elevated in ALS patients’ cerebrospinal fluid were tested for their effects on PD-L1 expression in human iPSC-derived motor neurons. Among the tested cytokines, only interferon-γ (IFN-γ) was found to strongly promote PD-L1 expression. Thus, we hypothesized that excessive exposure to IFN-γ may contribute to motor neuron degeneration in ALS. We observed that neuronal populations exposed to IFN-γ exhibited severe TDP-43 cytoplasmic aggregation and excitotoxic behavior correlated with impaired neural firing activity, hallmarks of ALS pathology, in both normal and ALS mutant (TARDB1K+/-) neurons. Single-cell RNA sequencing revealed possible mechanisms for these effects. Motor neurons exposed to IFN-γ exhibited an extensive shift of their gene expression profile toward a neurodegenerative phenotype. Notably, IFN-γ treatment induced aberrant expression levels for 70 genes that are listed in the recent literature as being dysregulated in various ALS subtypes. Additionally, we found that genes related to neuronal electrophysiology, protein aggregation, and TDP-43 misregulation were abnormally expressed in IFN-γ treated cells. Moreover, IFN-γ induced a significant reduction in the expression of genes that encode indispensable proteins for neuromuscular synapse development and maintenance, implying that the continuous cytokine exposure could directly impair signal transmission between motor axons and muscle membranes. Our findings suggest that IFN-γ could be a potent upstream pathogenic driver of ALS and provide potential candidates for future therapeutic targets to treat sporadic forms of ALS, which account for roughly 90% of reported cases.

## 1. Introduction

Amyotrophic lateral sclerosis (ALS) is a neurodegenerative disease that primarily attacks motor neurons, leading to progressive and ultimately fatal denervation of skeletal muscle throughout the body. Familial mutations are implicated as the cause for roughly 10% of ALS cases, while the pathologic origin of the other 90% remains unclear.^1, 2^ Drugs currently approved for use in ALS patients in the USA and/or Europe prolong patients’ lives by only a few months, leaving the disease intractable.^1^ Most patients do not develop neuromuscular symptoms until their 50s or 60s, regardless of the cause. However, the disease progresses rapidly once patients become symptomatic, with the average life expectancy of patients reported as less than 5 years after symptom onset.^1, 2^

A common pathologic feature of ALS is dysfunction of the RNA/DNA-binding protein, TDP-43. It has been suggested that TDP-43 plays a central, but poorly understood, role in ALS etiology since mutations of the gene encoding TDP-43, TAR DNA-binding protein (*TARDBP*), segregate with disease in some cases of familial ALS, and loss of nuclear TDP-43, associated with aberrant cytoplasmic accumulation of TDP-43 aggregates, is present in 97% of both familial and non-familial forms of ALS.^3–5^

Neuroinflammation is known to be a major factor in ALS progression and inflammatory cytokine action may switch from a neuroprotective to a neurotoxic role as the disease develops.^6–8^ Although previous studies have shown that activated peripheral immunity could acutely damage motor neurons’ survival and function, the molecular drivers of inflammatory stress and the following neurodegenerative consequences in neurons are unclear.^7, 9–12^ To assess the capacity of motor neurons to respond to various neuroinflammatory cytokines, we examined their effects on expression of Programmed death-ligand 1 (PD-L1), a cell surface protein, that is highly expressed by many cell types in response to exposure to a variety of inflammatory cytokines. PD-L1 plays a key role in immune escape by activating its cognate receptor, PD-1 on immune cells.^13^ We, therefore, investigated the influence of pro-inflammatory cytokines, known to be increased in ALS patients’ cerebrospinal fluid, on expression levels of PD-L1. We used both WT and ALS mutant (TARDBP^Q331K+/-^) iPSC-derived motor neurons for the cytokine screening assay, to compare their response.

We showed that interferon-γ (IFN-γ), was the only cytokine, among those tested, that strongly promotes neuronal PD-L1 expression in both groups. Although IFN-γ contributes to normal brain development and has been suggested to influence neurodevelopmental disorders such as autism and schizophrenia,^14^ a pathologic role for IFN-γ in upstream ALS etiology has not been widely explored. Importantly, we report that the prolonged IFN-γ exposure to human motor neurons in both normal and TARDBP^Q331K+/-^ groups recapitulated pathologic hallmarks of ALS, such as hyperexcitation followed by loss of neuronal firing ability and cytoplasmic aggregation of TDP-43.

In order to reveal mechanisms that might mediate the effects of IFN-γ exposure and TARDBP^Q331K+/-^ genotype on motor neuron ALS pathophysiology, we employed single-cell RNA sequencing to identify genes that are differentially expressed in response to IFN-γ exposure or TARDBP mutation, individually or in concert. The results of our transcriptomic analysis suggest that IFN-γ could be a potent neurodegenerative cue to induce ALS pathogenic processes and reveal possible mechanisms linking IFN-γ exposure to TDP-43 proteinopathy. Collectively, our findings suggest that compounds capable of inhibiting IFN-γ-mediated signaling in neurons may constitute promising targets for future immunotherapeutic drugs to prevent the progression of neurodegeneration in both familial and sporadic forms of ALS.

## 2. Results

### 2.1 IFN-γ significantly elevates PD-L1 expression on iPSC-derived motor neurons

We established a differentiation protocol, using small molecules, for generating induced pluripotent stem cell (iPSC)-derived ventrospinal motor neuron populations, essentially as described by Smith *et al*.^15, 16^ **(Supplementary Figure S1)** Furthermore, we previously employed CRISPR-Cas9-mediated gene editing techniques to introduce a pathogenic Q331K mutation into the TARDBP gene locus of the WTC11 iPSC line, creating an ALS (TDP-43 mutant) line and an isogenic control pair.^15^ In this study, these iPSCs were induced to generate human spinal motor neurons with wildtype (WT) and TARDBP^Q331K+/-^ genotypes. We assessed whether exposure of neurons with these two genotypes to inflammatory cytokines affected expression of PD-L1. Neurons were exposed to IFNα, IFN-γ, IL-4, IL-6, IL-1β, IL-17, or TNFα for 3 days (post-neuronal induction), which are the inflammatory cytokines reported to be most significantly increased in ALS patients.^17, 18^ (**Figure 1A, 1B**) Of the 7 listed cytokines, only IFN-γ promoted a significant increase in PD-L1 expression, and this was consistent across both WT and Q331K+/- populations. (**Figure 1C**)

**Figure 1.**
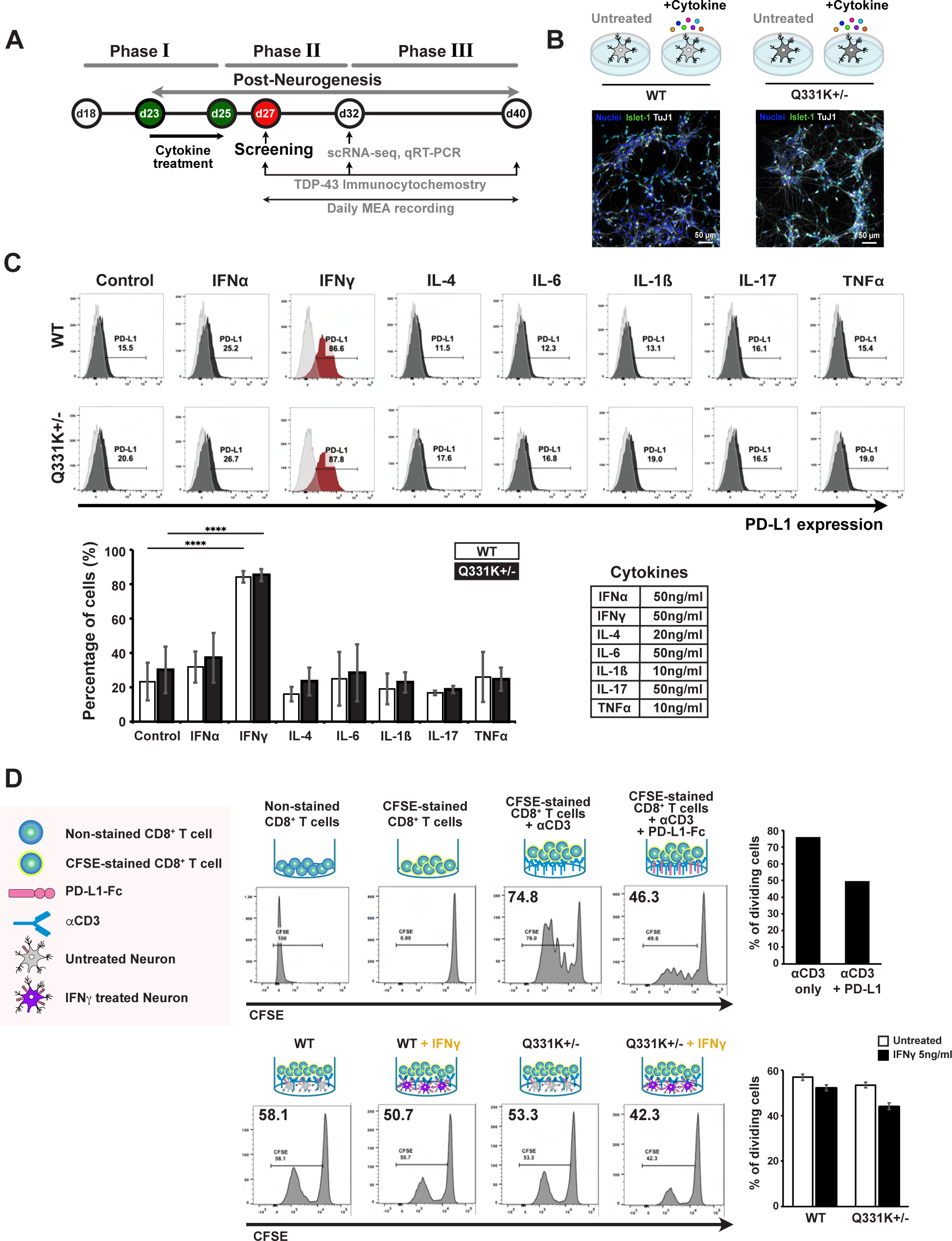
IFN-γ elevates PD-L1 expression in both WT and TARDBP mutant iPSC-derived spinal motor neurons. (A) Experiment timeline for motor neuron differentiation and downstream assays. (B) Schematic illustration of 4 experimental groups and culture images of WT and mutant neurons before cytokine treatment. (C) Cell surface PD-L1 expression after IFNα, IFNγ, IL-4, IL-6, IL-1β, IL-17, and TNFα treatment in iPSC-derived cells during phase II. Representative flow cytometry histograms (top) and quantification of flow cytometry analyses (bottom left). Error bars, S.D. (n=3, biological replicates). The table shows the concentrations of the treated cytokines (bottom right). (D) Suppression of CD8^+^ T cell proliferation by co-culture with IFN-γ-treated neurons. Schematic of the cells and antibodies used (left panel). CFSE-labeled CD8^+^ T cells were stimulated by incubating them in anti-CD3-coated plates for 5 days, in the absence or presence of IFN-γ-treated iPSC-derived neurons. Quantification of CFSE-labeled CD8^+^ T cell proliferation is depicted in each histogram (middle panels) and summarized in the bar graph (right panel).

Does PD-L1 in motor neurons function as a suppressor of cytotoxic T cell attack as it does in many cancer cell types? To address this question, we isolated T cells from a normal donor’s blood sample and labeled them with a carboxyfluorescein succinimidyl ester (CFSE), which covalently attaches a fluorescent label to the amines of cellular proteins. IFN-γ treated neurons were co-cultured with CFSE labeled CD8^+^ T cells for 5 days in the presence of anti-CD3, a marker of T cell proliferation. Labeled T cells without neurons used as controls in the presence of anti-CD3 or in the presence of both anti-CD3 and PD-L1-Fc proteins. PD-L1-Fc inhibited the proliferation of CD8^+^ T cells by approximately 30%. (**Figure 1D**) As shown in the same figure, IFN-γ treated neurons also reduced CD8^+^ T cell proliferation but only by approximately 10%, suggesting that the level of PD-L1 expression in motor neuron achieved by IFN-γ exposure is only modestly immunosuppressive.

### 2.2. IFN-γ exposure hyperactivates neurons followed by a complete loss of their firing function

Having verified that motor neurons are particularly sensitive to IFN-γ, we investigated whether this inflammatory cytokine might contribute to the electrophysiological dysfunction of motor neurons associated with ALS. We assessed function of iPSC-derived motor neuron cultures, with both WT and TARDBP^Q331K+/-^ genotypes, via multi-electrode arrays (MEAs), which enable recording of spontaneous field potential changes from cultured neurons in a non-destructive and non-invasive manner. (**Figure 2A**) We treated neurons daily with 3 different concentrations of IFN-γ (0.5 ng/mL, 5 ng/mL, 50 ng/mL) from days 23 to 40 post neuronal induction while recording the neuron’s firing activity across the same culture period. (**Figure 2B**) The time-course analysis of the mean firing rate shows a distinct difference in the firing pattern between untreated and IFN-γ-treated neurons. Untreated neurons showed a gradual increase in the firing rate as they matured for both the normal and mutant groups and maintained firing rates above 5 Hz from day 27 until the end of phase 2. However, neurons treated with IFN-γ above 5 ng/mL showed a drastic increase in the firing rate from day 27, reaching higher than 30 Hz by day 32, followed by a sharp decline in the firing rate subsequently. As a consequence of IFN-γ exposure, both WT and mutant groups completely lost their firing ability by day 38. (**Figure 2C**)

**Figure 2.**
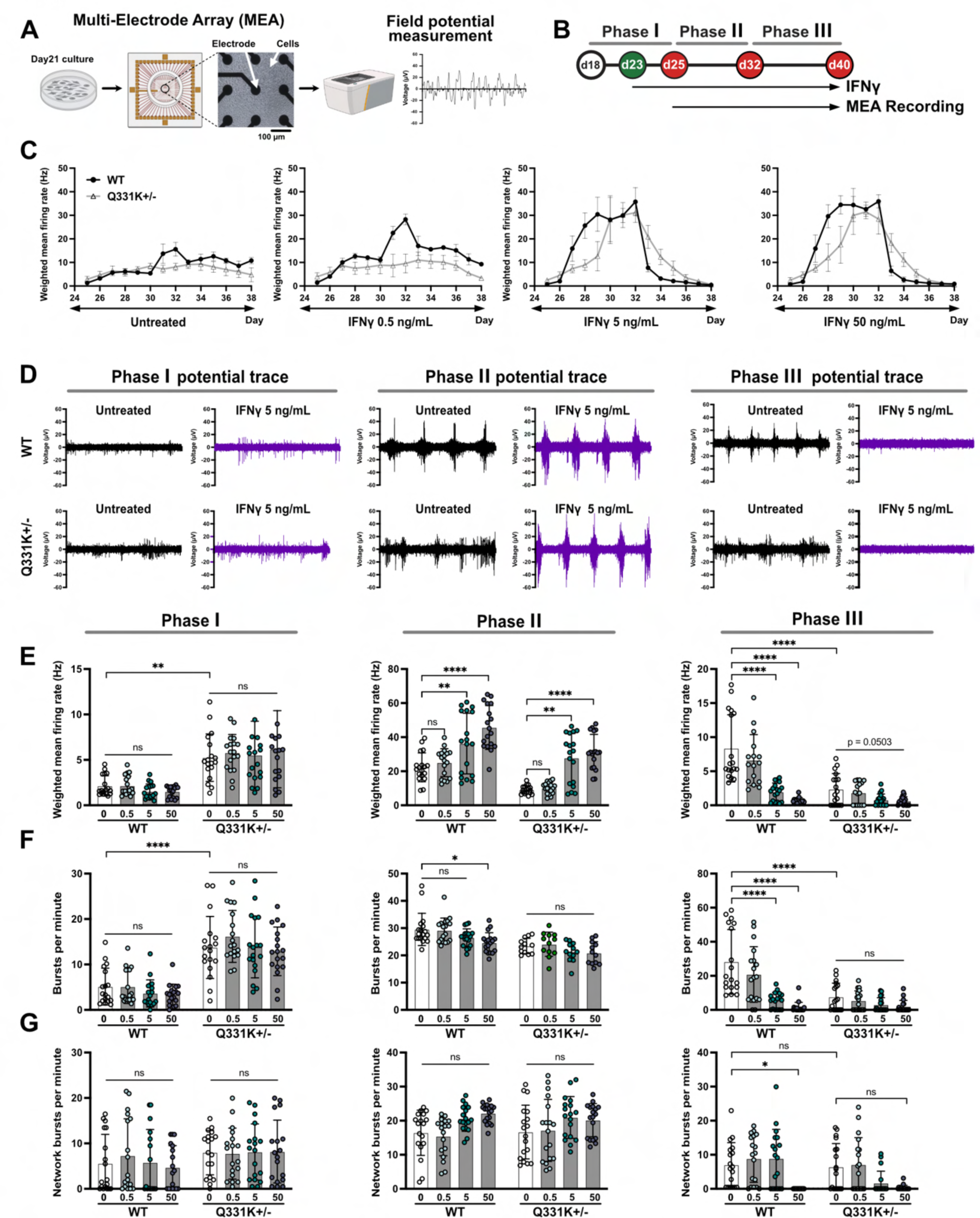
IFN-γ treatment induces aberrant electrophysiological activity in WT and Q331K+/- mutant iPSC-derived neuronal populations. (A) Illustration of the MEA experiment. (B) Experiment time line for the IFN-γ treatment and neuronal electrophysiology recordings. (C) Representative plot for time-course weighted mean firing frequency of the neurons. High concentration IFN-γ treated neurons show aberrant increase in firing rate during phase II, followed by an acute silencing in phase III. (D) Field potential trace of the neurons from each condition showing aberrant electrophysiological progression in response to IFN-γ treatment. Collective summary of phase-dependent electrophysiology of neurons for (E) weighted mean firing rate, (F) burst firing frequency, and (G) network burst firing frequency under the different stimulation conditions and mutation statuses. (Error bars S.D., n = 3 biological replicates; *P < 0.05, ***P<0.001, ****P<0.0001, ns=not significant; two-way ANOVA)

Although IFN-γ treatment was the major driver of this aberrant firing behavior, we also observed a notable difference in firing patterns between IFN-γ untreated WT and mutant neurons. The Q331K+/- mutant neurons tended to exhibit more active firing patterns than WT neurons early in the recording period. However, they started to exhibit impaired firing ability from day 31 and this trend was maintained until the end of the recording period. Notably, the firing frequency of mutant neurons was significantly lower than that of WT neurons at day 38. This result suggests that electrophysiological impairment could be one of the downstream pathologic effects of TARDBP mutation in neurons, which agrees with previous studies.^15, 19^ (**Figure 2C**) However, since IFN-γ exposure induced a much more pronounced impairment of neuronal electrophysiology, regardless of mutation status, the data suggest that IFN-γ-mediated inflammatory stress in the neurons may be a potent trigger of sporadic form of ALS. (**Figure 2D**)

This “hyperactivation followed by silencing” pattern was consistently observed in both WT and mutant neurons exposed to IFN-γ above 5 ng/ml. The weighted mean firing rate consistently showed a significant increase in firing frequency followed by a sudden drop, whereas untreated controls were still able to fire in the last phase of recording. (**Figure 2E**) In neuronal cultures, neural circuit formation can lead to synchronization of neural activity, detected as “burst” firing on MEAs.^16, 20, 21^ The differentiation protocol employed for these studies specifies neurons with ventral spinal cord identity, including interneurons as well as motor neurons, and the presence of interneurons was confirmed by our single-cell RNAseq analysis (data not shown). Consequently, these cultures have the potential to generate neuronal networks *in vitro*. We therefore monitored network burst firing behavior in these cultures to understand whether IFN-γ treatment altered inter-neuronal signaling. We observed an increase in the burst firing frequency of both WT and mutant neurons during the mid-phase, but the contribution of IFN-γ to this change was limited. However, the burst firing ability of mutant neurons was dramatically reduced in the last phase, regardless of IFN-γ treatment. IFN-γ -treated WT neurons showed dose-dependent decreases in burst firing ability in this same phase. (**Figure 2F**) However, we observed an impaired network burst firing frequency only in WT neurons under prolonged exposure to the IFN-γ, implicating the cytokine produced evidence of decaying inter-neuronal firing activity more prominently in the normal group of neurons. (**Figure 2G**)

Taken together, we observed that electrophysiological function of cultured neurons is affected by both TARDBP mutation and IFN-γ-mediated stress. However, the physiologic influence of IFN-γ treatment surpassed the inherent mutation in terms of the strength of its effect on firing patterns. We found that normal neurons were similarly vulnerable to continuous IFN-γ-mediated inflammatory stress, strongly suggesting that IFN-γ could be a critical pathogenic driver of sporadic ALS, which accounts for the majority of patient cases.

### 2.3. Prolonged IFN-γ exposure triggers cytoplasmic TDP-43 mis-localization

Does IFN-γ exposure contribute to triggering the cytoplasmic mis-localization of TDP-43, which is a defining hallmark of ALS? To address this question, we treated both WT and mutant neurons with 5 ng/mL IFN-γ for 15 days following the neurogenic phase of differentiation and immunostained the culture with anti-TDP-43 on days 25, 32, and 40 to monitor its expression and subcellular localization. (**Figure 3A**) Interestingly, we did not observe cytoplasmic TDP-43 mis-localization at any examined time-point in either WT or Q331K+/- mutant neurons without IFN-γ treatment. However, IFN-γ exposure led to prominent cytoplasmic accumulation and aggregation of TDP-43, in both groups of neurons. This phenotype was not observed until day 40, suggesting that TDP-43 mis-localization may require an accumulation of inflammatory stress over time to trigger the phenotype. (**Figure 3B**) Although the WT neurons developed a significant TDP-43 proteinopathy in response to IFN-γ exposure, the degree of the phenotype was more severe for mutant cells. This result indicates that the TARDBP mutation generates neurons that are more susceptible to cytoplasmic TDP-43 aggregation in response to inflammatory stress. (**Figure 3C**)

**Figure 3.**
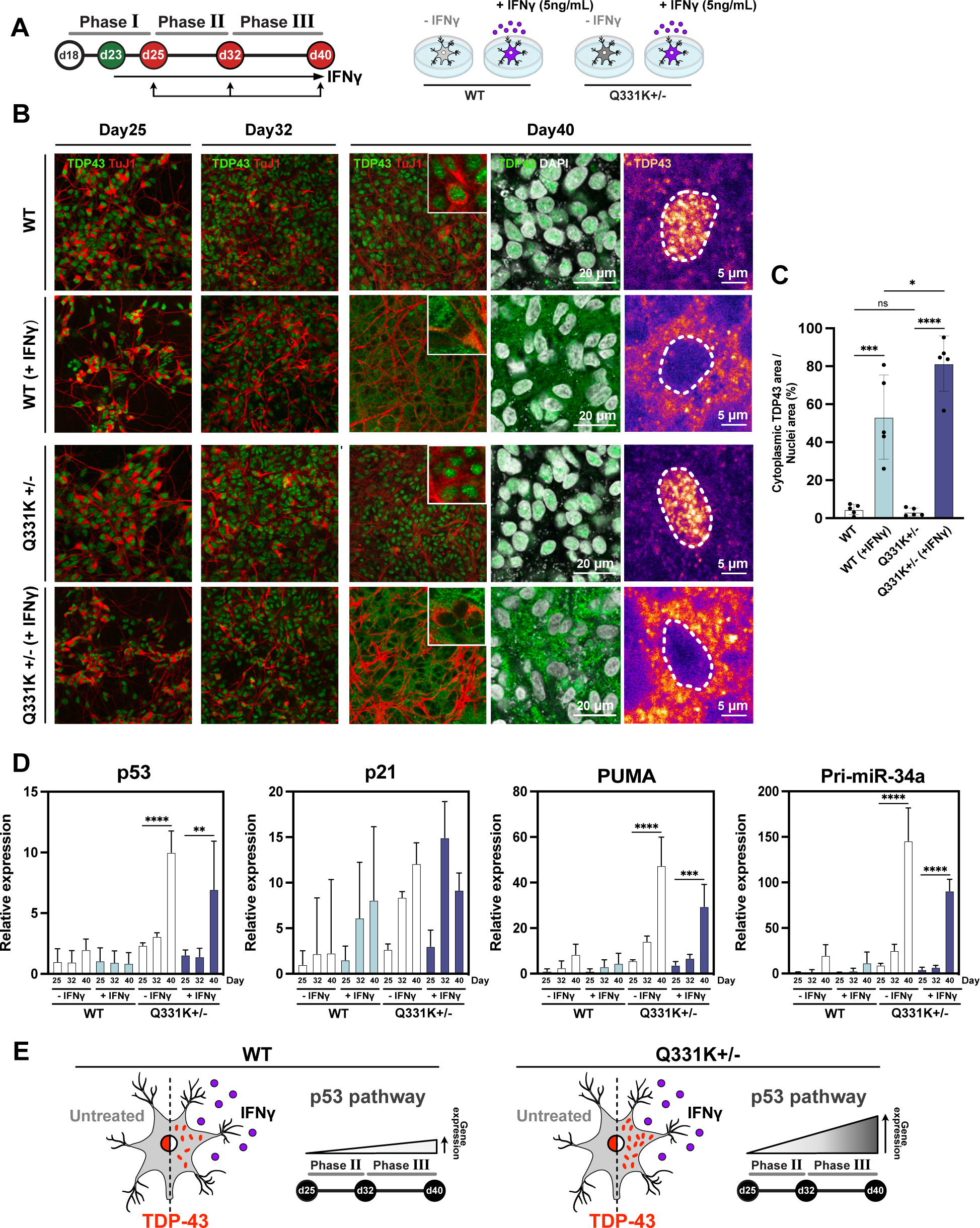
IFN-γ treatment triggers TDP-43 proteinopathy, whereas TARDBP mutation increases the basal activity of the p53 signaling pathway. (A) Experiment timeline of the immunocytochemistry and gene expression assays. (B) Immunocytochemistry assay of WT and Q331K+/- mutant cultures showing significant cytoplasmic translocation and aggregation of TDP-43 protein in response to IFN-γ stimulation at the later phases of both cultures. (C) Quantitative analysis of cytoplasmic TDP-43 translocation for each experimental condition. Time-course qRT-PCR analysis for each experimental group to quantify the expression of (D) p53, p21 (the early transcriptional target of p53 signaling), PUMA (a proapoptotic gene induced by p53), and Pri-miR-34a (a p53-regulated microRNA). An apparent trend in p53 pathway activation is observed with TARDBP mutation, whereas IFN-γ stimulation plays a limited role on this pathway. (E) Schematic summary of the phenotypic observation in each condition. (Error bars S.D., n = 3 biological replicates; *P < 0.05, ***P<0.001, ****P<0.0001, ns=not significant; one-way ANOVA)

### 2.4. Impact of IFN-γ exposure on the p53 pathway

TDP-43 promotes the function of p53, a key mediator of cell death across many diseases. Maor-Nof *et al*. recently reported that p53 could be a critical regulator of neurodegeneration in *C9orf72* mutant ALS, which is the most common subtype of familial ALS and is known to promote TDP-43 proteinopathy.^22, 23^ Hence, we further investigated the activation of p53 under the same conditions as in Figure 3B. We employed qRT-PCR to examine the expression of mRNA encoding p53, as well as its downstream effectors, p21, PUMA, and pri-miR-34a. (**Figure 3D**) Interestingly, the expression of those p53 pathway genes except p21 in cells bearing mutant TDP-43 all increased in a time-dependent manner, regardless of IFN-γ exposure, indicating that mutant TDP-43 may activate the p53 pathway through PUMA and Pri-miR-34a, even in the absence of extensive cytoplasmic TDP-43 mis-localization. Since p53 pathway activation is a common response to various types of cellular stress, this result suggests that the presence of mutant TDP-43 could drive an intracellular stress response to turn on p53 signaling in neurons, whereas IFN-γ-mediated inflammatory stress elicits significant TDP-43 proteinopathy regardless of the p53 activation. (**Figure 3E**)

### 2.5. Exposure to IFN-γ significantly alters the transcriptomic profile of both WT and TARDBP mutant motor neurons

Functional effects of IFN-γ are primarily mediated by JAK1/2 dependent activation of STAT1/2 transcription factors. Therefore, to gain insight into the mechanisms responsible for IFN-γ effects on electrophysiological dysfunction and TDP-43 cytoplasmic mis-localization, we characterized the transcriptional changes induced by exposure of motor neurons to IFN-γ. WT and Q331K+/- mutant neurons were treated with IFN-γ (5 ng/mL) for 10 days (days 22 to 32 post neuronal induction) directly following the neurogenic phase of differentiation, and subjected to single-cell RNA-sequencing on the last day of stimulation. We generated four single cell libraries defined by mutation status and IFN-γ treatment. (**Figure 4A**) Population analysis showed that motor neurons constituted 37.7% and 40.0% of WT and Q331K+/- mutant populations, respectively. The majority of non-neuronal cells in both groups were early glial progenitors and their gene expression profiles were all affected by IFN-γ treatment. (**Figure 4B, Supplementary Figure S2A, S2B)** Further cluster annotation, based on homeobox (HOX) gene expression patterns, indicated that both WT and Q331K+/- mutant motor neuron populations expressed regional identities specific to the upper spinal cord. **(Supplementary Figure S2C)**

**Figure 4.**
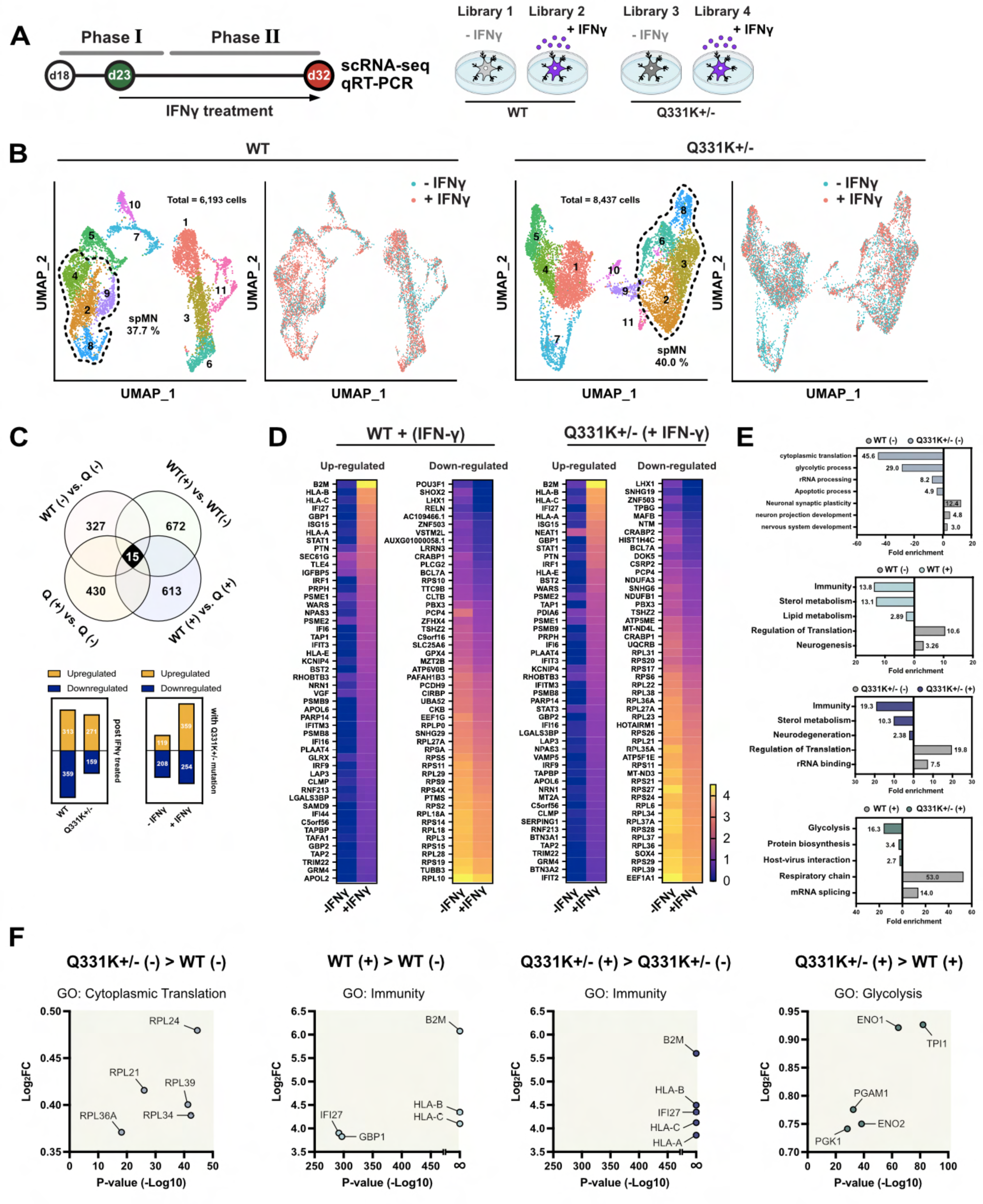
IFN-γ treatment induces inflammatory gene expression in both WT and Q331K+/- mutant iPSC-derived motor neurons. (A) Experiment timeline of IFN-γ treatment and gene expression analysis. The schematic illustration shows the library constitution for the single-cell RNA sequencing experiment. (B) UMAP clustering of both cultures shows the populational distribution of the differentiated cells. (C) The number of DEGs in each pairwise comparison. (+) indicates culture with IFN-γ treatment and (–) indicates untreated controls. Q indicates Q331K+/- mutant group. (D) Heatmap of top 50 DEGs based on the fold change post IFN-γ treatment in both WT and mutant motor neurons. Scale indicates relative gene expression level in each group. Gene counts per cell were normalized for their total counts over all genes and logarithmized. (E) Gene ontology (GO) analysis of the DEGs generated in each pairwise comparison. (F) The top 5 DEGs included in the selected GO terms in each pairwise comparison. > denotes the listed genes are more expressed in the left group than in the control group on the right.

Motor neuron stimulation by IFN-γ generated 672 (WT) and 430 (Q331K+/-) differentially expressed genes (DEGs) compared to untreated controls. Interestingly, 613 DEGs appeared between the IFN-γ stimulated WT and Q331K+/- motor neurons, implying a potential role for the TDP-43 mutation in mediating neuronal responses to IFN-γ. We also found 327 DEGs between the WT and Q331K+/- neurons that were not exposed to IFN-γ, indicating that the TDP-43 mutation exerted a significant impact on cultured motor neuron’s downstream gene expression profiles. (**Figure 4C**)

A heatmap of the top 50 DEGs in each group shows that IFN-γ exposure significantly increased inflammatory gene expression and decreased ribosomal protein gene expression in both WT and Q331K+/- motor neurons. A notable elevation of B2M (ß-2-microglobulin) expression was observed in both WT and Q331K+/- motor neurons in response to IFN-γ treatment. B2M is a component of MHC-1 (Major Histocompatibility Complex-I) class molecules. They are expressed in all nucleated cells and are responsible for presenting foreign peptide fragments on the cell surface, which bind CD8+ T cells to trigger an immediate immune response. ß-2-microglobulin is a critical mediator of MHC-1 surface expression, verified using a B2M-defienct mouse model that rarely presents MHC-1 on the cell surface and exhibits poor development of CD8+ T-cells.^24, 25^ The significant increase of B2M expression in our culture model demonstrated that IFN-γ elicits an acute inflammatory response from treated motor neurons. In addition, a sharp increase of HLA-A, HLA-B, and HLA-C expression in WT and Q331K+/- neurons, which encodes human leukocyte antigens that correspond to MHC-I class, confirmed that the IFN-γ-mediated inflammatory response constituted a major upstream driver of gene expression change in both groups of motor neurons. (**Figure 4D**)

Gene ontology analysis was performed using the DEGs that arose from 4 different pairs of gene expression data, to generally characterize the nature of the effects. Changes in gene expression due to the TARDBP mutation were mainly related to cytoplasmic translation, glycolytic processes, and neuronal synaptic plasticity. Within the cytoplasmic translation GO term, ribosomal proteins (including RPL24, RPL21, RPL39, RPL34, and RPL36A) were up-regulated in mutant neurons. These data mirror the sequencing results previously reported from primary excitatory neurons harboring a *C9orf72* mutation, which is responsible for the most frequent form of familial ALS.^26^ DEGs arising from IFN-γ treatment were primarily related to immunity, sterol metabolism, and regulation of translation in both WT and mutant neurons. However, mutant neurons exposed to IFN-γ exhibited significant differences in the expression levels of genes related to glycolysis, respiratory chain, and mRNA splicing, when compared to WT motor neurons receiving the same concentration of IFN-γ. (**Figure 4E, 4F)**

### 2.6. IFN-γ drives gene expression changes characteristic of neurodegeneration

We sought to ascertain whether exposure to IFN-γ promoted the expression of ALS-related transcripts in cultured motor neurons and, if so, how the trend differed from that caused by the TARDBP^Q331K+/-^ mutation. We compared the identified DEGs individually to lists of the ALS risk genes recently identified by applying a machine learning algorithm to ALS GWAS data and molecular profiling of iPSC-derived motor neurons.^27^ (**Figure 5A**) We found a significant overlap between the 4 sets of DEGs with those risk genes. In particular, we found 72 DEGs related to ALS pathology, derived by IFN-γ exposure. The result confirms that cytokine exposure puts cultured neurons on to a neurodegenerative pathway, which could potentially lead to the development of an ALS phenotype, although the IFN-γ did not induce motor neuron death in either WT or mutant groups. (**Figure 5B, 5C**)

**Figure 5.**
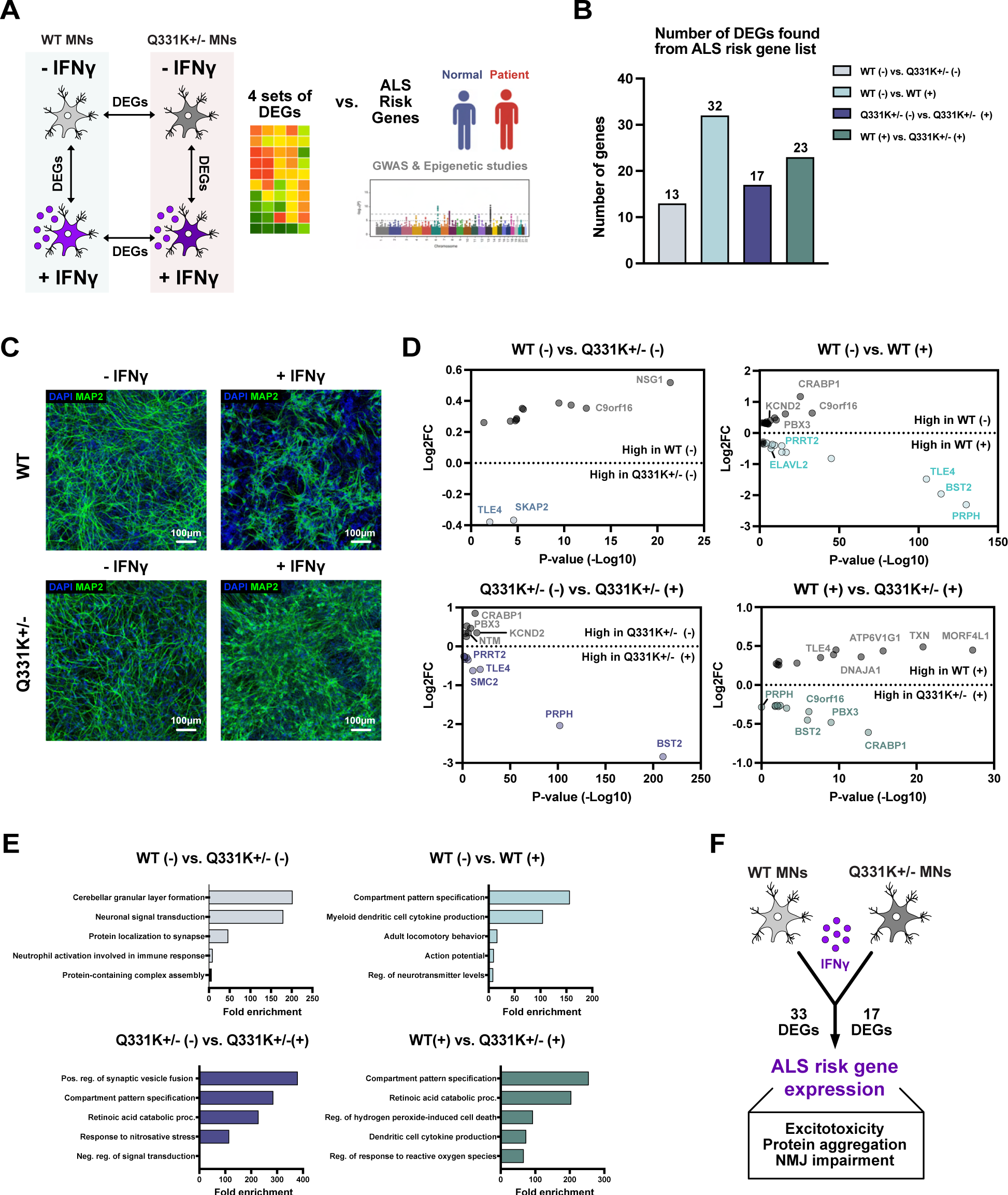
The IFN-γ-induced transcriptomic changes promote an ALS phenotype in motor neurons. (A) The schematic illustration shows how the analysis was performed. (B) DEGs found from each comparison were compared with the previously reported risk genes for various neurodegenerative disease states. The number of risk genes on the y axis shows how many of the reference risk genes for each disease were overlapped with the DEGs generated by either mutation or IFN-γ treatment. (C) The immunostaining images show that IFN-γ treatment does not induce notable cell death in both groups of culture. (D) The fold-change and p-value of selected DEGs for each comparison that have a significant role in neurodegeneration based on the recent literature. (E) GO analysis results for the subgroup of DEGs that are known to increase the risk of ALS development. (F) Illustration summary of the discovery delineated in the figure. (+) indicates culture with IFN-γ treatment and (–) indicates untreated controls.

The most significantly mis-regulated DEGs in each pairwise comparison, and the GO analysis results for those genes, revealed a distinct effect of Q331K mutation and IFN-γ exposure on motor neuron function, suggesting that the functional behavior of these cells is more greatly affected when both factors are present. (**Figure 5D, 5E**) In summary, IFN-γ treatment significantly altered the gene expression profile of both WT and Q331K+/- mutant motor neurons in a manner that could lead to the development of ALS phenotypes. (**Figure 5F**)

From the list of genes exhibiting greater than two-fold differential expression for each pairwise comparison, we employed qPCR to confirm and more precisely quantify the extent of differential expression of genes that are particularly relevant to neurodegeneration. We found that apoptotic genes, such as IFI27, IFIT2, and PLAAT4, were highly elevated in response to IFN-γ treatment but were unaffected by the Q331K mutation. (**Figure 6A**) We also confirmed decreasing CRABP1 gene expression following IFN-γ exposure. CRABP1 is an essential component of neuromuscular junction (NMJ) development and maintenance via the CRABP1-CaMKII-Agrn pathway. Its reduced expression in IFN-γ treated cells indicates that this cytokine may directly impair the stability and function of muscle innervation by motor neurons. (**Figure 6B**) Moreover, the list of DEGs in IFN-γ treated cells compared with controls included genes encoding ion channel proteins responsible for regulating neuronal electrophysiology. RNA-seq and qRT-PCR data both implicated that the genes for the voltage-gated potassium channel KCND2 and the sodium channel SCN9A were mis-regulated in a direction that might promote neuronal excitotoxicity. (**Figure 6C**) The significant increase in expression of PRPH that encodes the neuron-specific intermediate filament protein, peripherin, is interesting because over-expression of peripherin in motor neurons promotes neuronal cell death, and peripherin is a component of ubiquitinated inclusions and of axon spheroids in ALS.^28–30^ Given that the overexpression of WT peripherin causes protein aggregation and motor neuron degeneration in transgenic mice, our result implies that exposure to IFN-γ could drive adoption of similar phenotypes in human motor neurons. Similarly, the decrease of TPBG observed following cytokine treatment indicates that IFN-γ could increase the risk of protein aggregation since the ablation of TPBG is known to promote alpha-synuclein aggregation in a mouse model of neurodegeneration. (**Figure 6D**) Lastly, we found significantly altered expression for ALS risk genes under IFN-γ exposure, such as BST2, PBX3, and C9orf16, which have been reported to be mis-regulated in various familial ALS models although their pathologic function is yet to be identified. (**Figure 6E**) In summary, IFN-γ exposure promoted gene expression changes that are likely to induce electrophysiological impairment, neuromuscular synapse breakdown, protein aggregation, and apoptotic cell death. (**Figure 6F**)

**Figure 6.**
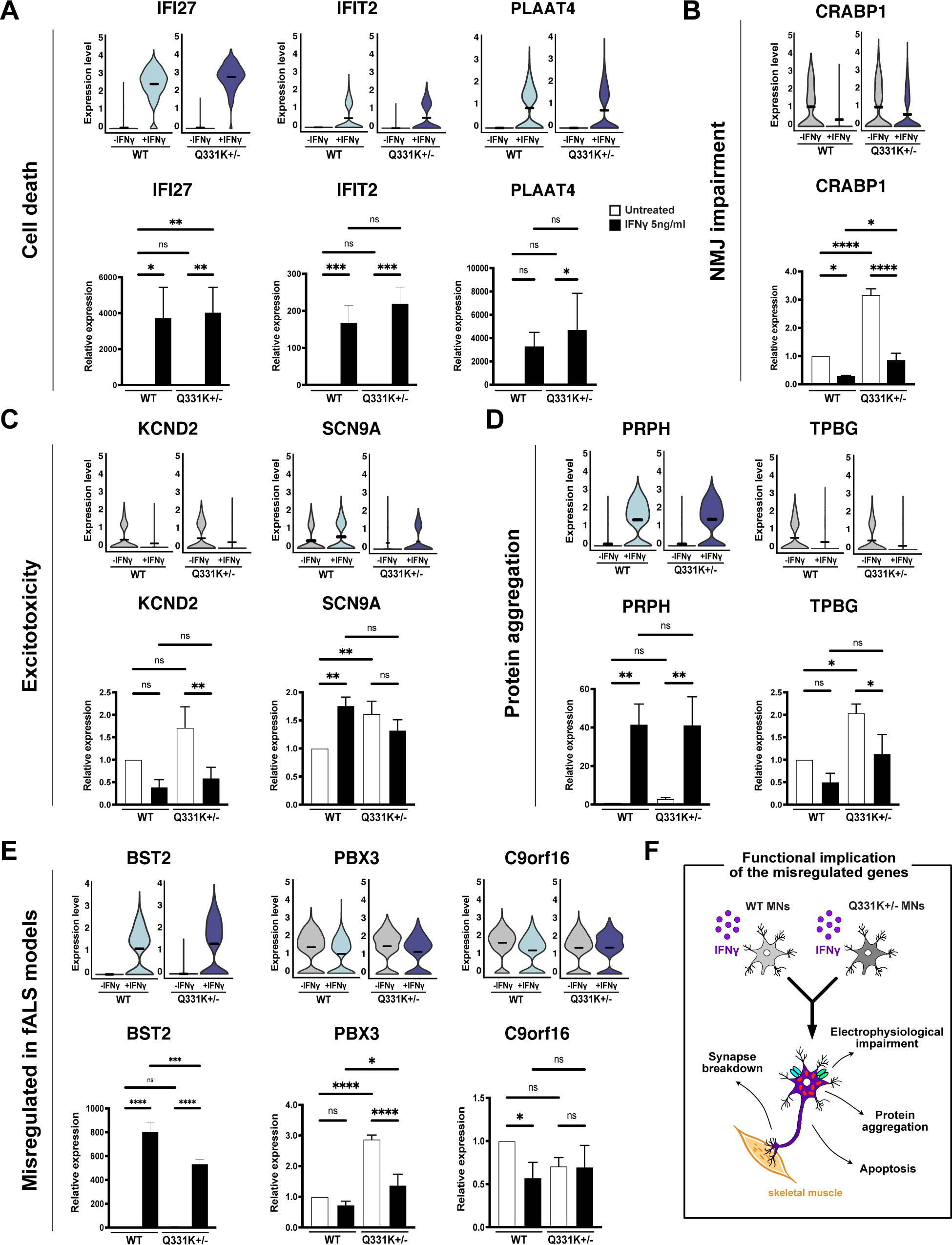
qRT-PCR verification of the selected DEG expression categorized by their functional implications in motor neurons. Comparison of single-cell RNAseq and qRT-PCR assay results for the DEGs potentially responsible for cell death (A), NMJ impairment (B), excitotoxicity (C), and protein aggregation (D). (E) Additional DEGs known to be mis-regulated in familial ALS models, in which aberrant expression was caused by IFN-γ treatment. (F) Schematic summary of the potential implications for the listed DEGs driving neurodegenerative phenotypes. (Error bars S.D., n = 3 biological replicates; *P < 0.05, ***P<0.001, ****P<0.0001, ns=not significant; two-way ANOVA)

## 3. Discussion

In this study, we found that only IFN-γ, among cytokines tested, increases the expression of immune-modulatory protein PD-L1 in both WT and Q331K+/- mutant motor neurons. Other pro-inflammatory cytokines known to be elevated in ALS patient cerebrospinal fluid were not able to induce the PD-L1 up-regulation, our data suggest that motor neurons might be particularly sensitive to IFN-γ. We hypothesized that an inflammatory response leading to excessive exposure of spinal cord motor neurons to IFN-γ could elicit expression of a neurodegenerative phenotype. The source of neuroinflammation-associated IFN-γ that may affect motor neurons remains to be fully charactered. T cells are a major source of IFN-γ secretion, although NK cells, glia and even neurons can produce IFN-γ.^31^ T cell infiltration of the CNS parenchyma occurs only in a few types of acute inflammation. A T cell population resides constitutively associated with the choroid plexus, where it may secrete IFN-γ into the CSF. Importantly, CSF of ALS patients exhibits an expansion of a T cell subpopulation expressing eomesodermin, which drives IFN-γ expression.^32^ IFN-γ is permeable to the blood spinal barrier, particularly in the cervical region, implying that T cells impacted by peripheral inflammation might expose motor neurons to IFN-γ.^33–35^ Moreover, Bonney *et al* showed that IFN-γ even promotes its permeability to the blood-brain barrier, implying that uncontrolled IFN-γ secretion may broadly affect the physiology of the entire CNS.^36^ Finally, peripherally projecting motor axons lie outside the blood brain barrier. Exposure of sympathetic axons to IFN-γ mediates axonal STAT1 phosphorylation and transport of these activated transcription factors to the somatic nucleus.^37^ If a similar retrograde IFN-γ axonal signaling pathway exists in motor neurons, T cell-generated IFN-γ circulating peripherally might affect motor neurons.

We examined the capacity of IFN-γ exposure to elicit two hallmarks of ALS – impaired neurophysiological function of motor neurons and mis-localization of TDP-43 in cytoplasmic aggregates, and observed the occurrence of both, in agreement with our hypothesis. We found that the electrophysiological function of neurons bearing a TARDBP mutation and exposed to IFN-γ was significantly impaired, and the impact of the cytokine treatment surpassed the effect of the mutation. The cytokine-treated neurons showed a ‘hyperexcitation and silencing’ pattern (regardless of whether or not they expressed a TARDBP mutation) that resembled the excitotoxic behavior of neurons in the mid-phase of ALS. Although further follow-up studies are required, downregulated genes encoding the potassium channels and abnormal elevation of genes for sodium channel proteins could be responsible for their functional damage.

We further studied whether TDP-43 proteinopathy, another pathologic hallmark of ALS, could be induced by IFN-γ treatment. The continuous treatment of IFN-γ triggered a severe cytoplasmic aggregation of TDP-43 in both WT and mutant neurons at late stages of culture. No such phenotype was induced at any time point by the presence of the TARDBP mutation alone, although the severity of the IFN-γ triggered phenotype was more pronounced in the mutant groups. This result implies that mis-localization of TDP-43 requires an upstream trigger beyond mutation in the TARDBP gene, but does not discount the possibility that the mutation in TARDBP does contribute to the development of a pathologic phenotype.

Our single-cell RNA sequencing analysis revealed that IFN-γ triggers a significant transcriptomic alteration in both WT and Q331K+/- mutant neurons that trends toward the development of an ALS phenotype and suggests several specific mechanisms by which increased IFN-γ transcriptional signaling in motor neurons might impair electrophysiological function and promote TDP-43 mis-localization. Among the DEGs arising from IFN-γ treatment, we identified genes known to be aberrantly expressed in familial ALS, as well as potential drivers of cell apoptosis, protein aggregation, excitotoxicity, and NMJ breakdown, implying that cytokine exposure may induce a broad range of ALS pathological hallmarks, even in motor neurons that do not bear mutations in TDP-43. Since many DEGs identified in familial ALS mouse models were also mis-regulated in our normal motor neurons after simply treating with IFN-γ, we could postulate that the mutation may work as a potential upstream trigger to induce secretion of IFN-γ, which in turn drives neurodegenerative gene expression in motor neurons.

We found that IFN-γ induces an abnormal expression of motor neuron-specific genes responsible for regulating NMJ development and maintenance. CRABP1 is a retinoic acid-binding protein, that was recently found to regulate motor function via maintenance of the NMJ through the CRABP-CaMKII-Agrin axis. Lin *et al*. showed that CRABP1 knock-out mice exhibited adult-onset ALS-like phenotypes and developed significantly impaired NMJs followed by motor axon degeneration, which was rescued after the re-expression of the gene.^38^ It is notable that, in our hands, CRABP1 expression was dramatically reduced in both genotypes of motor neurons under exposure to IFN-γ, indicating that excessive immunological activity might contribute to the development of sporadic forms of ALS via direct targeting of the neuromuscular synapse. PRPH encodes a neuronal intermediate filament protein, peripherin. Mutations in this gene lead to the enrichment of one of the three splice variants observed in human patients (Per28), which causes a disruption of neurofilament assembly and is associated with the development of an ALS phenotype.^28^ Peripherin levels are significantly elevated in patients with motor neuron disease and the protein accumulates in the majority of axonal inclusion bodies in ALS patient motor neurons. ^28–30, 39^ Given that motor neurons are highly polarized cells with extremely long axons, the regulation of cytoskeletal genes is of critical importance in the axonal trafficking of organelles and biomolecules, which directly affects motor neuron survival and function. PRPH also localizes to Bunina bodies, which are intraneuronal protein inclusions, suggesting that an increase in peripherin expression may cause protein aggregation in motor neurons.^28^ Xiao *et al*. showed that overexpression of PRPH induces self-aggregation and causes motor neuron degeneration in transgenic mice. In our experiments, the significant increase of PRPH in motor neurons exposed to IFN-γ raises two important questions: 1) Does increased PRPH in neurons disrupt axonal trafficking and degenerate neurons in a retrograde manner? 2) Does PRPH also induce cytoplasmic aggregation of TDP-43? TPBG encodes trophoblast glycoprotein and a deficiency in this protein induces the aggregation of alpha-synuclein in mice. Although TPBG mis-regulation is more directly linked to Parkinson’s disease, the downstream effect of decreased TPBG expression in motor neurons may be worth investigating in order to establish a better understanding of its potential role in developing a broad range of neurodegenerative phenotypes.

Regardless, there still are important questions that to be answered to better understand the pathologic role of TDP-43 proteinopathy on ALS development. First, more studies are required to understand the pathologic consequence of severe TDP-43 cytoplasmic aggregations induced by IFN-γ in motor neurons. Second, speculative mechanisms mentioned here linking particular DEGs to IFN-γ-mediated protein aggregation need to be experimentally tested. Finally, we still do not understand the exact role of TARDBP mutations in aggravating the disease phenotype once initiated by cytokine exposure.

We additionally investigated whether the p53 pathway, a cell death mediator gene that is known to be mis-regulated in familial ALS is turned on by IFN-γ treatment. Given that p53 is one of TARDBP’s downstream target genes, we postulated that the p53 pathway could be initiated by IFN-γ-induced TDP-43 proteinopathy. However, IFN-γ stress did not induce p53 signaling. Instead, the presence of a TARDBP mutation strongly upregulated p53 pathway genes in a time-dependent manner, indicating that the p53 pathway is directly affected by the mutation status of TDP-43.

We believe these results raise important questions, the answers to which could lead to a better understanding of the pathogenic role of aberrant immunity-mediated neurodegeneration, which could be a key component driving rapid disease progression in ALS. For example, it is currently unclear whether the inflammatory reaction occurs at the axon terminal, the periphery of the patient’s body, before spreading the pathologic effect to neuronal cell bodies in a retrograde manner. Finally, it would be valuable to investigate the neuron-autonomous pathologic role of PD-L1, besides its immune-modulatory function, since recent studies have revealed that exogenous PD-L1 treatment to mouse sensory neurons significantly impairs their firing ability without having any influence on the immune response.^40^ Therefore, understanding the neuron-autonomous role and downstream signaling path of PD-L1-mediated neuromodulation could be a critical step towards understanding the mechanism of the IFN-γ-induced neurodegeneration.

## 4. Conclusion

Our findings strongly suggest that IFN-γ is a potent driver of neuroinflammation that induces the adoption of ALS phenotypes in both WT and Q331K+/- mutant spinal motor neurons. We found that IFN-γ treatment not only induces a significant increase in functional PD-L1 and inflammatory gene expression but also extensively shifts motor neuron’s gene expression towards a diverse array of neurodegenerative features. We found that continuous exposure to IFN-γ alters motor neuron phenotypes in a manner that mirrors changes observed in ALS. Both WT and TARDBP mutant iPSC-derived motor neurons showed a dramatic decay of firing frequency coupled with an increase in cytoplasmic TDP-43 aggregation following IFN-γ treatment. However, the mutant neurons exhibited more severe phenotypes for both features, suggesting a role for the mutation in exacerbating the inflammatory stress response. We further confirmed that IFN-γ exposure triggered transcriptomic alterations that suggest movement towards expression of a neurodegenerative phenotype in motor neurons. Thus, our study provides strong evidence for a pathogenic link between IFN-γ-mediated inflammation and subsequent neurodegeneration in spinal motor neurons that mirrors ALS pathology. These results suggest that immunomodulatory compounds may constitute strong targets for ameliorating symptoms in both sporadic and familial forms of ALS.

## 5. Materials and Methods

### Maintenance of human iPSC lines

WTC11 (WT, Q331K+/-) iPSCs were frozen in mFreSR medium (Stem Cell Technologies) and stored under cryogenic conditions in liquid nitrogen. 10 cm dishes were treated with Matrigel (Thermo Fisher Scientific) diluted 1 to 60 in DMEM/F12 medium (Thermo Fisher Scientific) and incubated at 37°C/5% CO2 overnight. On the day of plating, vials of cells were removed from liquid nitrogen storage and incubated in a 37°C water bath for 3 minutes. The content of the vial (1 mL) was then transferred to a 15 mL centrifuge tube and centrifuged for 3 minutes at 300 g. The supernatant was aspirated and cells were resuspended in fresh mTeSR (Stem Cell Technologies) supplemented with 10 µM Y-27632 (Thermo Fisher Scientific), a specific inhibitor of Rho kinase activity. Prepared Matrigel plates were then aspirated, washed with PBS containing Ca^2+^ and Mg^2+^ (Thermo Fisher Scientific), and the cell suspension was then plated evenly over the prepared culture surface. Y-27632 was removed from the culture medium the first day after plating and cells were fed daily with fresh mTeSR from then on. Cells were incubated at 37°C/5% CO_2_ until iPSC colonies filled the field of view when visualized using an Eclipse TS100 microscope (Nikon) fitted with a 10X lens. At this point, medium was aspirated and replaced with TrypLE 1× solution (Invitrogen), incubated in 37°C/5% CO_2_ for 5 minutes. The detached cell colonies were dissociated with gentle trituration with P1000 pipette, followed by adding the same volume of PBS with calcium and magnesium. The cell suspension was spun down and resuspended in fresh mTeSR with mild trituration to gently break up cell clusters. The cell suspension was then split across the desired number of Matrigel-coated plates and returned to the incubator. During continued culture, any iPSC colonies displaying irregular boundaries, significant space between cells or low nuclear to cytoplasmic ratios were carefully marked and removed from culture using a fire-polished, sterile glass pipette.

### Differentiation of iPSCs into spinal motor neurons

Human iPSCs were passaged onto Matrigel-coated 6-well plates as described above and incubated at 37°C/5% CO_2_ in mTeSR until they reached ∼50% confluency. At this point, cultures were differentiated into regionally unspecified neural progenitor cells using a monolayer differentiation method. Briefly, the undifferentiated day0 cells were treated with dual-SMAD inhibitors, SB431532 and LDN193189, as well as ascorbic acid (AA). CHIR99021 was added to the medium for the purpose of activating the Wnt pathway, which has been proven to enhance neuroepithelial differentiation and proliferation. At day 5, cultures were treated with all-trans retinoic acid (RA) for caudalization and the SHH agonist Purmorphamine, to give ventralization cues to the cells. On day 11, cells were passaged onto fresh Matrigel coated 6-well plates and treated with RA and AA until day 18 to expand the neural progenitor population. These cells were then passaged at 100,000 cells/cm^2^ onto 0.01% poly-L-ornithine (Sigma-Aldrich) and 5 µg/mL laminin (Sigma-Aldrich)-coated surfaces and exposed to culture conditions promoting motor neuron differentiation. Specifically, cells were fed with 10 ng/mL of neurotrophic factors (BDNF, GDNF, IGF-1 from R&D systems, NT3 from Stem Cell Technologies), ascorbic acid (2.27 µM). During all stages of differentiation, a basal medium consisting of a 1:1 mix of Neurobasal medium and DMEM/F12, supplemented with B27, Glutamax, N2, Non-essential amino acids, and penicillin-streptomycin, was used. All cells used in the described experiments were differentiated from WTC11 colonies between passages 45 and 55.

### Cytokine treatment

WT and Q331K+/- mutant iPSC-derived neurons were exposed to 7 different cytokines, treated from day 23 to day 25 of the ventrospinal neuron differentiation method described above. Recombinant human and mouse IFNα, IFNγ, IL-4, IL-6, IL-1, IL-17, and TNF-α were used at concentrations depicted in Figure 1B. All of the cytokines were purchased from R&D Systems.

### Co-culture of CD8+ T cells and iPSC-derived motor neurons

Whole blood was collected into 10 mL BD Vacutainer Plastic Collection Tubes with Sodium Heparin and inverted multiple times. To isolate PBMCs, whole blood was gently layered over a double volume of Ficoll in a Falcon tube and centrifuged for 30-40 minutes at 400g without a brake. Four layers formed, each containing different cell types - the uppermost layer contained plasma, which was removed by pipetting. The second layer contained PBMCs and these cells were gently removed using a pipette and added to PBS to wash off any remaining platelets. The pelleted cells were then counted and the percentage viability estimated using Trypan blue staining. A 96 well cell culture plate was coated with a CD3-specific Ab (OKT3, eBioscience for human T cells) and 5 μg/mL concentrations of PD-L1-Ig for control experiments. For co-culture of CD8^+^ T cells and iPS derived motor neurons, the culture plate was coated with poly-L-ornithine, laminin, and a CD3-specific Ab sequentially. Human CD8^+^ T cells were purified from PBMCs freshly isolated from whole blood using CD8a^+^ T cell isolation kit II, human (Miltenyi Biotec.). Purity was confirmed to be over 90 % by flow cytometry. The T cells were labeled with 1μM CFSE, quenched by cold FBS, and incubated in plates coated with CD3-specific Ab. Human iPSC-derived neurons were cocultured with CD8^+^ T cells purified from PBMCs at a 1:1 ratio (Neurons : CD8^+^ T cells) for 5 days in culture medium with CD3 stimulation. On day 5 after stimulation, cells were stained with an Ab against CD8-APC and then analyzed for CFSE dilution by flow cytometry.

### Flow cytometry

The following antibodies were used for flow cytometry analysis: PD-L1 (29E.2A3, BioLegend), CD44 (C44Mab-5, BioLegend), and CD8 (SK1, BioLegend). The stained cells were analyzed using a BD FACS Calibur flow cytometer. For each sample, 10,000 events were recorded and histograms were plotted. The percentages of cells were calculated using FlowJo software (Treestar, Ashland, OR).

### Immunocytochemistry

Cells were fixed in 4% paraformaldehyde for 15 minutes and permeabilized in 0.2% Triton-x solution, followed by blocking with 5% goat serum in PBS for 1 hour at room temperature. Cells were then incubated with primary antibodies diluted in 0.5% BSA in PBS overnight at 4°C. The next day, cells were washed 3 times with PBS. They were then incubated in a secondary antibody solution containing secondary antibodies diluted in 0.5% BSA in PBS overnight at 4°C. Counterstaining was performed with Vectashield containing DAPI (Vector Labs). Images were taken at the Garvey Imaging Core at the University of Washington’s Institute for Stem Cell and Regenerative Medicine using a Leica SP8 Confocal System on an inverted microscope platform. 12-bit 2048×2048 pixel images were acquired with Leica LAS X software. Antibodies used in this study were as follows: mouse anti-Islet1 (1 in 200, DSHB), rabbit anti-β III tubulin (1 in 500, Sigma Aldrich), rabbit anti-TDP43 (1 in 200, Invitrogen), goat anti-ChAT (1 in 200, R&D Systems), Alexafluor-488 conjugated goat-anti-mouse secondary antibody (1:500, Invitrogen), Alexafluor-594 conjugated goat-anti-rabbit secondary antibody (1:500, Invitrogen) and Alexafluor-647 conjugated Donkey-anti-goat secondary antibody (1:500, Invitrogen).

### Electrophysiology

Population level function in motor neuron cultures was assessed using 48-well multielectrode arrays (MEA) in conjunction with the Maestro MEA system (Axion Biosystems). Day 21 cells were plated on Cytoview 48 well plates (Axion Biosystems) at the density of 100,000 cells per well. Plated cells were then subjected to daily cytokine and blocking antibody treatment using fresh medium from the next day onwards. Electrophysiological recordings were taken every day from day 22 to day 40 for all experimental groups. During data acquisition, standard recording settings for spontaneous neuronal spikes were used (Axis software, version 2.5), and cells were maintained at 37°C/5% CO_2_ throughout the 2-minute recording period. The standard settings have 130 × gain, and record from 1 to 25 000 Hz, with a low-pass digital filter of 2 kHz for noise reduction. In all experiments, spike detection was set at 5× the standard deviation of the noise and network burst detection was recorded if at least 25% of the electrodes in a given well showed synchronous activity. Reported results were calculated by averaging all of the electrodes in each well, then averaging data from duplicate wells.

### Quantitative real-time PCR

We extracted total RNA from cells using Trizol (Invitrogen). After checking RNA purity and concentration, RNA was reverse transcribed to cDNA using the iScript SuperMix reagent (BioRad). Primers (IDT) were diluted in nuclease-free water with PowerUp SYBR Green master mix (Applied Biosystems) and qPCR performed on the Applied Biosystems 7300 machine. The following primers were used human β-actin: forward, 5’ -ACTCTTCCAGCCTTCCTTCC- 3’, reverse, 5’ -CAATGCCAGGGTACATGGTG- 3’; human PD-L1: forward, 5’ -GGAAATTCCGGCAGTGTACC-3’, reverse, 5’ -GAAACCTCCAGGAAGCCTCT- 3’; human IFI27: forward, 5’ – TGGAATGCCACGGAATTAACC - 3’, reverse, 5’ - GCCACAACTCCTCCAATCAC - 3’; human IFIT2: forward, 5’-CACTGCAACCATGAGTGAGA- 3’, reverse, 5’ – AGGTTGCACATTGTGGCTTT - 3’; human PLAAT4: forward, 5’ – CCGCTGTAAACAGGTGGAAA - 3’, reverse, 5’ - CCGCTGTAAACAGGTGGAAA - 3’; human CRABP1: forward, 5’ - AAGGTCGGAGAAGGCTTTGA- 3’, reverse, 5’ - AGTGGCTAAACTCCTGCACT- 3’; human KCND2: forward, 5’- CCTTCTTCTGCTTGGACACG-3’, reverse, 5’- TCTGTCATCACCAGCCCAAT-3’; human SCN9A: forward, 5’- AGGACCTCAGAGCTTTGTCC-3’, reverse, 5’- AAGTCACTGCTTGGCTTTGG-3’; human PRPH: forward, 5’- CTCAAGCAGAGGTTGGAGGA-3’, reverse, 5’- CGTGCAGCTTCTTGAGGAAC-3’; human TPBG: forward, 5’ - GCCTCGTCCAACAACCATAC- 3’, reverse, 5’ - CACCTCTTCGCCTCTTGTTG- 3’; human BST2: forward, 5’ - CCATCTCCTGCAACAAGAGC-3’, reverse, 5’ – TGCATCCAGGGAAGCCATTA - 3’; human PBX3: forward, 5’- TTACCAAGGGTCCCAAGTCG-3’, reverse, 5’- GTAGCCTCCCGTCTGATTGA-3’; human C9orf16: forward, 5’- GCAGAATACGCTGCCATCAA-3’, reverse, 5’- CGTGGAGGTGGTCATTCTTC-3’; human p53: forward, 5’- CCCAAGCAATGGATGATTTGA-3’, reverse, 5’- GGCATTCTGGGAGCTTCATCT-3’; human p21: forward, 5’- GGCAGACCAGCATGACAGATT-3’, reverse, 5’- GCGGATTAGGGCTTCCTCT-3’; human PUMA: forward, 5’- CCTGGAGGGTCCTGTACAATCT-3’, reverse, 5’-GCACCTAATTGGGCTCCATCT-3’; human pri-miR-34a: forward, 5’- CCAGAACAGTTCCTGCTGC-3’, reverse, 5’- TTAGCTGGTGCTCTCAGAC-3’. Relative RNA levels were calculated from Ct values.

### Single cell RNA sequencing

Single cells differentiated from WT and Q331K+/- mutant iPSC-derived neurons with or without IFNγ treatment were collected on day 32 of differentiation, to generate total 4 independent single cell libraries. Following single cell dissociation using TrypLE (Thermo Fisher Scientific), cells were filtered through a 70 µm filter and spun down at 1200 rpm for 5 minutes. The single cell pellet was resuspended in neuron maintenance medium at a maximum concentration of 2,000,000 cells/mL, followed by measuring cell viability using the Countess II automated cell counter (Invitrogen), which confirmed over 91% viability of cells in all groups. Cells were spun down at 1200 rpm for 5 minutes again and resuspended in PBS + 0.04% BSA at a concentration of 15,000 cells/16.5 µL to target harvesting 10,000 recovered cells per library. cDNAs of single cells from each group were barcoded with 10× Genomic’s Chromium Next GEM Chip G Single Cell Kit and Chromium Next GEM Single Cell 3’ Kit v3.1, by following manufacturer’s protocol. Constructed libraries were loaded to the Tapestation (Agilent) to quantify the cDNA amount and confirm fragmentation status, followed by sequencing with the Illumina NextSeq2000 at the Genomics core with the University of Washington’s Institute for Stem Cell and Regenerative Medicine. The sequencing was performed at an estimated read depth of 10,000 reads/cell. Sequenced FASTQ reads were initially processed using CellRanger v.3 software with settings recommended by the manufacturer (10X Genomics).

### Transcriptomic data analysis

Read alignment, filtering, barcode counting, and unique molecular identifier (UMI) counting was performed using CellRanger data processing software provided by 10× Genomics. The feature barcoded matrix was obtained for further gene expression analysis with Seurat V4.0. The quantitative summary of sequencing was obtained from the CellRanger as follows: 1) The sequencing of single cells from WT culture without IFNγ stimulation detected 1,911 genes per cell, 4,105 UMI per cell, and 21,312 mean reads per cell. 2) The sequencing of single cells from WT culture with IFNγ stimulation detected 1,833 genes per cell, 3,726 UMI per cell, and 16,427 mean reads per cell. 3) The sequencing of single cells from Q331K+/- mutant culture without IFNγ treatment detected 1,651 genes per cell, 3,443 UMI per cell, and 18,452 mean reads per cell. 4) The sequencing of single cells from Q331K+/- mutant culture with IFNγ treatment detected 1,858 genes per cell, 3,803 UMI per cell, and 20,319 mean reads per cell. For the comparative gene expression analysis, the integration of single-cell data described by Stuart *et al*. was applied on the basis of the Seurat algorithm. Before applying the integration, cells with low UMI counts, doublets, and cells with relatively high mitochondrial DNA content were removed. The cleared datasets were subsequently normalized in order to correct for differences in read depth and library size, using Seurat’s “LogNormalize” function which divides feature counts of the cell by its total counts, followed by multiplying the scale factor. Then, the integration method included in Seurat v.4 aligned the shared cell populations across multiple datasets to identify cells that are in a matched biological state in each library, which allows comparative gene expression analysis across the selected libraries to be performed. The PCA was performed with the integrated data post normalization, followed by running the “RunUMAP” function to generate 2-dimensional UMAP projections using the top principal components detected in the dataset. “FindConveredMarkers” function in Seurat identified conserved cell type markers for generating the feature plots with the “FeaturePlot” function. The clusters were annotated by analyzing the list of conserved marker genes in each cluster with the gene ontology database, The Human Protein Atlas, and the PanglaoDB. Differential gene analysis was performed by using the “FindMarkers” function in Seurat, and the result was displayed with volcano plots, heatmaps, and violin plots to visually highlight the contrast in gene expression across the selected transcriptomic datasets.

### Statistical data analysis

All experiments were performed at least in triplicate, and repeated using 2-3 independent differentiation runs for WTC11 neurons except for the single cell RNA sequencing. Significant differences between groups were evaluated using unpaired t-tests for two conditions, or one-way ANOVA, with *post hoc* tests for multiple comparisons, for experiments with 3 or more groups. Mann Whitney U tests and ANOVA on ranks were used to analyze the statistical significance of differences arising between sets of non-normally distributed data. In all experiments, a p value of less than 0.05 was considered significant. All statistical tests were performed using the GraphPad Prism statistics software.

## Acknowledgements

This work was funded by NIH R03 TR004009 (Alec S.T. Smith) and supported by a Predoctoral fellowship (Changho Chun) from the Institute for Stem Cell and Regenerative Medicine (ISCRM) at the University of Washington. Additional funding was provided by a Sponsored Research Agreement from Curi Bio Inc. and a philanthropic gift from the Eileen & Larry Tietze Foundation (both awarded to David L. Mack), and by an Elsa U Pardee Foundation (Jung Hyun Lee). The single-cell RNA sequencing work performed in this study was supported by the ISCRM Genomics Core. Our thanks to Dr. Mary C. Regier for her assistance with the single-cell library preparation for the sequencing experiment and subsequent data analysis.

## Author contribution

Conceptualization: CC, JHL

Methodology: CC, JHL

Investigation: CC, JHL

Visualization: CC, JHL

Funding acquisition: ASTS, DLM, JHL

Project administration: CC, JHL

Supervision: ASTS, MB, DLM

Writing – original draft: CC, JHL

Writing – review & editings: ASTS, MB, DLM

