## Supplementary Figure S1, S2 for "Interferon-γ Elicits Pathological Hallmarks of ALS in Human Motor Neurons"

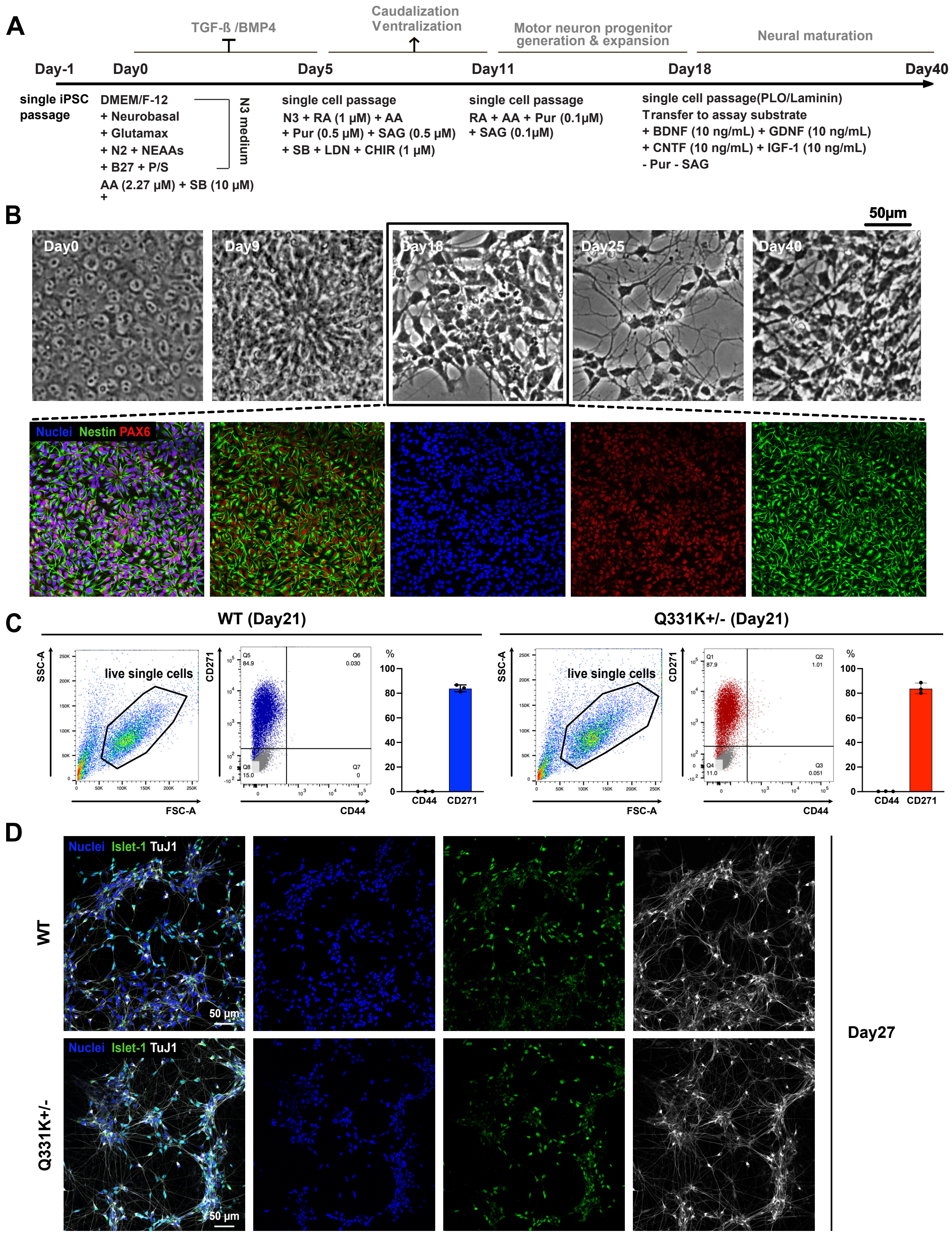

**Figure S1. Motor neurons are differentiated from both WT and Q331K+/- mutant iPSCs**. (A) Differentiation scheme of iPSC-derived motor neurons. (B) Morphological change of cells during the differentiation. Day 18 cultures highly express Nestin and PAX6, demonstrating a neural progenitor phenotype. (C) Flow cytometry analysis with day 25 cultures shows minimal inclusion of non-neuronal cells (CD44+) and enrichment of neurons (CD271+). (D) Immunocytochemistry assay of day 27 cultured neurons with the pan-neuronal marker (TuJ1) and motor neuron-specific marker (ISLET-1), demonstrating successful motor neuron differentiation.

### Supplementary Figure S2

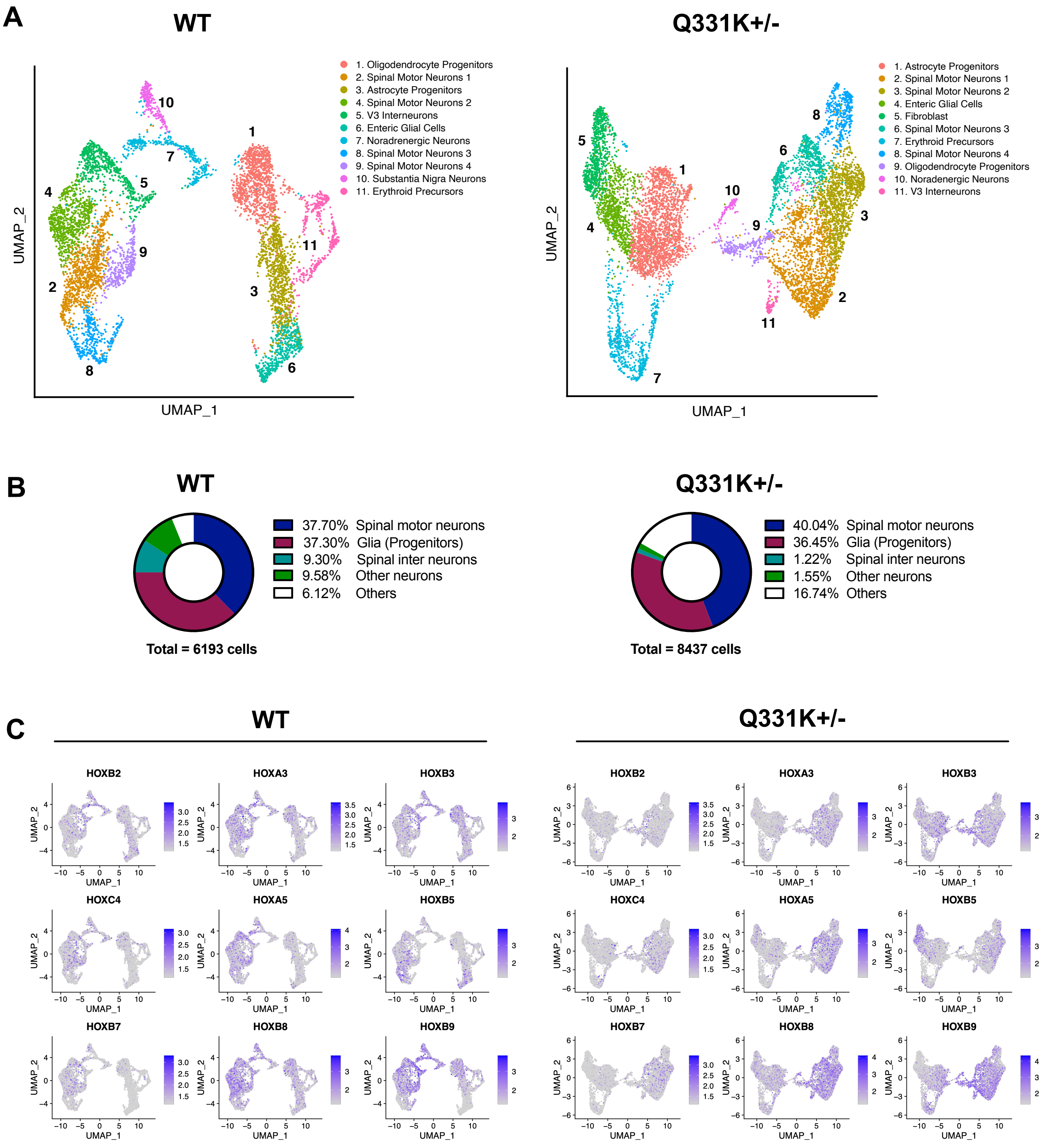

**Figure S2. Population analysis of differentiated culture from WT and Q331K+/- mutant iPSCs**. (A) UMAP of day 32 culture differentiated from WT and Q331K+/- mutant iPSCs without IFN**γ** stimulation. Identity of 11 clusters in each UMAP was annotated on the top right side of each panel. (B) Quantitative analysis of cell population classified to 5 groups. Both cultures are enriched with spinal motor neurons and glial progenitors. (C) Feature plot of HOX gene expression for all of the single cells in both WT and Q331K+/- mutant culture.
